# Improved draft reference genome for the Glassy-winged Sharpshooter (*Homalodisca vitripennis*), a vector for Pierce’s disease

**DOI:** 10.1101/2021.06.04.447158

**Authors:** Cassandra L. Ettinger, Frank J. Byrne, Mathew A. Collin, Derreck Carter-House, Linda L. Walling, Peter W. Atkinson, Rick A. Redak, Jason E. Stajich

## Abstract

*Homalodisca vitripennis* (Hemiptera: Cicadellidae), known as the glassy-winged sharpshooter, is a xylem feeding leafhopper and an important agricultural pest as a vector of *Xylella fastidiosa,* which causes Pierce’s disease in grapes and a variety of other scorch diseases. The current *H. vitripennis* reference genome from the Baylor College of Medicine’s i5k pilot project is a 1.4-Gb assembly with 110,000 scaffolds, which still has significant gaps making identification of genes difficult. To improve on this effort, we used a combination of Oxford Nanopore long-read sequencing technology combined with Illumina sequencing reads to generate a better assembly and first-pass annotation of the whole genome sequence of a wild-caught Californian (Tulare County) individual of *H. vitripennis*. The improved reference genome assembly for *H. vitripennis* is 1.93-Gb in length (21,254 scaffolds, N50 = 650 Mb, BUSCO completeness = 94.3%), with 33.06% of the genome masked as repetitive. In total, 108,762 gene models were predicted including 98,296 protein-coding genes and 10,466 tRNA genes. As an additional community resource, we identified 27 orthologous candidate genes of interest for future experimental work including phenotypic marker genes like *white*. Further, as part of the assembly process, we generated four endosymbiont metagenome-assembled genomes (MAGs), including a high-quality near complete 1.7-Mb *Wolbachia* sp. genome (1 scaffold, CheckM completeness = 99.4%). The improved genome assembly and annotation for *H. vitripennis*, curated set of candidate genes, and endosymbiont MAGs will be invaluable resources for future research of *H. vitripennis*.

## Introduction

*Homalodisca vitripennis*, commonly known as the glassy-winged sharpshooter, is a xylem-feeding leafhopper, non-model insect in the order Hemiptera and an important agricultural pest of grapes, citrus, and almonds (Turner and Pollard 1959; Blua *et al.* 1999). The full native range of *H. vitripennis* includes the southeastern USA and northeastern Mexico (Triapitsyn and Phillips 2000). However, since its invasion into California in the 1990s, it has proliferated to be the most extensive vector in California of *Xylella fastidiosa*, the causative agent of Pierce’s disease (Sorensen *et al.* 1996; Redak *et al.* 2004; Stenger *et al.* 2010; Backus *et al.* 2012). Unfortunately, the long-term use of insecticides to control *H. vitripennis* has led to high levels of resistance in California populations (Byrne and Redak 2021).

Although both a transcriptome and draft genome for *H. vitripennis* are available, we believe there is value in expanding and improving on these resources (Nandety *et al.* 2013; Hunter *et al.* 2016). The current *H. vitripennis* reference genome (Hvit v.2.0) from the Baylor College of Medicine’s i5k pilot project is a 1.4-Gb assembly with 110,000 scaffolds from a lab-reared Florida line. The assembly still has significant gaps making identification of genes difficult. Likely contributing to this is the large size and repetitive nature of many insect genomes (Cernilogar *et al.* 2011; Jiang *et al.* 2012); for example, repetitive regions make up to 40% of the genomes of silkworms (Cai *et al.* 2012), 47% in mosquitos (Nene *et al.* 2007) and 60% in locusts (Wang *et al.* 2014). The use of long-read sequencing can improve genome contiguity when repetitive regions are present (Richards and Murali 2015). In addition to improving genome contiguity for annotation purposes, an improved assembly would enable the ability to look into chromosomal-level rearrangements, like those observed in other Hemiptera to occur as a selection for insecticide resistance (Manicardi *et al.* 2015).

Using a combination of Oxford Nanopore long-read sequencing technology combined with Illumina-sequencing reads, we report an improved assembly of the *H.* v*itripennis* genome and genome annotation. We briefly describe the repetitive-sequence landscape of the *H.* v*itripennis* genome and identify candidate genes of interest for future experimental work. Finally, we identify and report on obligate and facultative endosymbiont genomes from the assembly. An improved genome for *H. vitripennis,* particularly from an invasive Californian individual, is a critical resource needed to support on-going management strategies (e.g. RNAi, CRISPR technologies, viral, etc), and studies of *H. vitripennis* population structure, which may be important for understanding resistance to non-biological controls.

## Methods & Materials

### Organism collection and sequencing

In August 2019, sharpshooters were collected from citrus groves across multiple locations in California as part of a study on imidacloprid resistance (Byrne and Redak 2021). Of these, three sharpshooters (designated A6, A7, A9) collected from an organic citrus grove (*Citrus sinensis* (L.) Osbeck) in Porterville, California (Tulare County) were used for genome sequencing. The insects from this location (Tulare-Organic) were confirmed to be susceptible to imidacloprid using a topical application bioassay (Byrne and Redak 2021).

Total DNA was extracted from three Tulare-Organic individuals (A6, A7, A9) following the 10X Genomics protocol for high molecular weight genomic DNA extraction from single insects (“DNA extraction from single insects” 2018). DNA from A6 was then constructed into a paired-end DNA library at UC Riverside Institute of Integrative Genome Biology (IIGB) Genomics Core and sequenced on an Illumina NovaSeq 6000 at the Vincent J. Coates Genomics Sequencing Laboratory at the University of California, Berkeley producing 97 Gb in 322 M Illumina reads. Additionally, DNA from all three Tulare-Organic individuals (A6, A7, A9) was sequenced on an Oxford Nanopore MinION using an R9.4.1 flow cell. Long-fragment DNA was validated using gel electrophoresis and Qubit (Invitrogen, Carlsbad, CA, United States). A total of 1.5 ug of high-quality DNA was prepared in singleplex with a SQK LSK-109 kit using End Prep, DNA Repair, and Blunt Ligase (New England Biolabs, Ipswich, MA) according to the Nanopore recommended protocol. Sequence reads were basecalled using Guppy version 3.3.0 on NVIDIA Tesla-P100 GPU in the UCR High Performance Computing Cluster (https://hpcc.ucr.edu).

Additional sharpshooters collected from California citrus groves in Porterville (Tulare-Organic), Temecula (Temecula-Organic), Bakersfield (GBR-Organic) and Terra Bella (Tulare-Conventional) were confirmed to have varying levels of imidacloprid resistance (Byrne and Redak 2021). Four sharpshooters were sampled from each of these locations for a total of 16 individuals that were processed for transcriptome sequencing (Byrne and Redak 2021). For each sharpshooter, RNA was extracted from adult prothoracic leg tissue tissue using Monarch Total RNA Mini Kit (New England Biolabs, Ipswich, MA). Paired-end RNA-Seq libraries were constructed with NEBNext Ultra II Directional RNA prep (New England Biolabs, Ipswich, MA) and sequenced on NovaSeq 6000 to produce an average of 87 M paired reads per library (minimum library 51 M, max library 124 M reads).

### Genome assembly

Genome assembly was performed with the susceptible (Tulare-Organic) individuals by sequencing A6 Illumina library and the A6, A7, and A9 Nanopore libraries. The assembler MaSuRCA v. 3.3.8 (Zimin *et al.* 2013), which performs read correction and extension was used in combination with Flye v. 2.5 (Lin *et al.* 2016; Kolmogorov *et al.* 2019) as implemented in MaSuRCA with parameters (“LHE_COVERAGE=35 LIMIT_JUMP_COVERAGE = 300 EXTEND_JUMP_READS=0 cgwErrorRate=0.20). Additional assembly parameters and related scripts, as well as all code used throughout this work, are available on GitHub and archived in Zenodo (Ettinger and Stajich 2021).

The resulting contigs were scaffolded against the existing reference assembly from the Baylor College of Medicine’s i5k pilot project (hereafter referred to as i5k) (i5K Consortium 2013; Hunter *et al.* 2016) available in GenBank (<u>GCA_000696855.2</u>) using Ragtag v. 1.0.0 (Alonge *et al.* 2019). Vector and contaminant screening were performed using the vecscreen option in AAFTF v0.2.4 (Stajich and Palmer 2019). Mitochondrial and endosymbiont genome identification and removal were performed as described in detail below. Assembly evaluation and comparison were performed using QUAST v. 5.0.0 (Gurevich *et al.* 2013) and BUSCO v. 5.0.0 (Simão *et al.* 2015) against both the eukaryote_odb10 and hemiptera_odb10 datasets. Assembly statistics and BUSCO status were visualized in R v. 4.0.3 using the tidyverse v. 1.3.0 package (Wickham *et al.* 2019; R Core Team 2020).

To investigate genome size and potential heterozygosity, we used jellyfish v. 2.3.0 (Marçais and Kingsford 2011) to count a range of *k*-mers (k = 19, 21, 23, 25, 27) and produce *k*-mer frequency histograms. We then supplied these histograms to GenomeScope v. 2.0 (Ranallo-Benavidez *et al.* 2020) and findGSE (Sun *et al.* 2018), which both provide estimates of genome size, percent heterozygosity and percent repeat content.

### Mitochondria and endosymbiont identification

The mitochondrial genome was assembled and identified from the Illumina reads using the “all” module in MitoZ v. 2.4-alpha (Meng *et al.* 2019). We then used Minimap v.2.1 (Li 2018) to map the mitochondrial genome against the draft *H*. *vitripennis* genome. Partial matches to the mitochondrial genome found in the draft *H*. *vitripennis* genome were subsequently hard masked. Mitochondria annotation was performed with MITOS2 (Donath *et al.* 2019) and the tbl file manually checked for gene name consistency and flagged discrepancies before conversion to sqn file format for upload to NCBI.

We used the BlobTools2 pipeline (Challis *et al.* 2020) to identify and flag scaffolds of microbial origin for possible removal. Taxonomy of each scaffold was putatively assigned using both diamond (v. 2.0.4) and command-line BLAST v. 2.2.30+ against the UniProt Reference Proteomes database (v. 2020_10) (Camacho *et al.* 2009; Buchfink *et al.* 2015; Boutet *et al.* 2016). We estimated coverage by mapping reads to the scaffolds with bwa (Li and Durbin 2009) and merged and sorted the alignments using samtools v. 1.11 (Li *et al.* 2009). We then used the BlobToolKit Viewer to visualize the resulting putative assignments.

As an alternative method to identify possible microbial contamination in the assembly, we ran the anvi’o v.7 pipeline (Eren *et al.* 2015). This involved first obtaining coverage information by mapping reads to scaffolds with bowtie2 v. 2.4.2 (Langmead and Salzberg 2012) and samtools v. 1.11 (Li *et al.* 2009). We then generated a scaffold database from the draft *H*. *vitripennis* genome using ‘anvi-gen-contigs-database’, which calls open-reading frames using Prodigal v. 2.6.3 (Hyatt *et al.* 2010). Single-copy bacterial (Lee 2019), archaeal (Lee 2019), and protista (Delmont) genes were then identified using HMMER v. 3.2.1 (Eddy 2011) and ribosomal RNA genes were identified using barrnap (Seemann). Putative taxonomy was assigned to gene calls using Kaiju v. 1.7.2 (Menzel *et al.* 2016) with the NCBI BLAST non-redundant protein database nr including fungi and microbial eukaryotes v. 2020-05-25. Next, an anvi’o profile was constructed for contigs >2.5 kbp using ‘anvi-profile’ with the ‘--cluster-contigs’ option, which hierarchically clusters scaffolds based on their tetra-nucleotide frequencies. Scaffolds were manually clustered into metagenome-assembled genomes (MAGs) using a combination of hierarchical clustering, taxonomic identity, and GC content using both ‘anvi-interactive’ and ‘anvi-refine’. MAG completeness and contamination were assessed using ‘anvi-summarize’ and then again using the CheckM v. 1.1.3 lineage-specific workflow (Parks *et al.* 2015). MAGs were taxonomically identified using GTDBTk v.1.3.0 (Chaumeil *et al.* 2019), which places bins in the Genome Taxonomy Database phylogenetic tree and putatively assigns taxonomy based on ANI to reference genomes and tree topology. MAG placement was visualized in R v. 4.0.3 using the ggtree v. 2.2.4 and treeio v. 1.12.0 packages (R Core Team 2020; Wang *et al.* 2020; Yu 2020).

We took the resulting scaffolds from both approaches (e.g. scaffolds that were taxonomically flagged as containing bacteria, archaea, or viruses reads by BlobTools2 and all scaffolds assigned to MAGs through the anvi’o workflow) and assessed whether to remove the scaffolds from the assembly using JBrowse2 (Buels *et al.* 2016). To do this, we converted diamond and BLAST taxonomy file outputs, as well as the BUSCO v. 5.0.0 (Simão *et al.* 2015) matches to both the eukaryote_odb10 and hemiptera_odb10 datasets, into GFF formatted files to enable their import into JBrowse2. After manual assessment of these scaffolds via JBrowse2, we proceeded with conservatively removing from the draft *H. vitripennis* genome only those scaffolds that were assigned to MAGs. Additional symbiont and mitochondrial regions that were identified by NCBI’s Contamination Screen were subsequently removed during deposition.

### Repetitive element annotation

Prior to gene annotation, we used RepeatModeler v. 2.0.1 (Flynn *et al.* 2020) and RepeatMasker v. 4.1.1 (Smit *et al.* 2013-2015) to generate and soft mask predicted repetitive elements in the draft *H. vitripennis* genome (Table S1). To visualize the repeat landscape, we used the parseRM.pl script v. 5.8.2 (https://github.com/4ureliek/Parsing-RepeatMasker-Outputs) with the ‘−l’ option on the RepeatMasker output (Kapusta *et al.* 2017). The parseRM.pl script calculates the percent divergence from the consensus for each predicted repeat using the Kimura 2-Parameter distance while correcting for higher mutation rates at CpG sites. Percent divergence can be a proxy for repeat element age with older elements expected to have higher divergence due to expected accumulation of more nucleotide substitutions relative to younger elements. Here, we chose to group repeats into bins of 1% divergence. Repeat landscapes were visualized in R v. 4.0.3 using the tidyverse v. 1.3.0 package (Wickham *et al.* 2019; R Core Team 2020).

### Genome annotation

To identify protein-coding genes and tRNAs, we used the Funannotate pipeline v. 1.8.4 on the masked genome (Palmer and Stajich 2020). Briefly this involved first training the gene predictors on the RNAseq data using Trinity v. 2.11.0 and PASA v. 2.4.1 (Haas *et al.* 2003; Grabherr *et al.* 2011). Next, gene prediction was performed using a combination of software including Augustus v. 3.3.3, GeneMark-ETS v. 4.62, GlimmerHMM v. 3.0.4 and SNAP v 2013_11_29 (Korf 2004; Majoros *et al.* 2004; Stanke *et al.* 2006; Ter-Hovhannisyan *et al.* 2008). Consensus gene models were then produced using EVidenceModeler v. 1.1.1 (Haas *et al.* 2008) and tRNAs were predicted using tRNAscan-SE v. 1.3.1 (Lowe and Eddy 1997). Consensus gene models were then refined using the RNAseq training data from PASA, which includes untranslated region (UTR) prediction. Protein annotations were then putatively assigned for consensus gene models based on similarity to Pfam (Finn *et al.* 2014) and CAZyme domains (Lombard *et al.* 2014; Huang *et al.* 2018) using HMMER v.3 (Eddy 2011) and similarity to MEROPS (Rawlings *et al.* 2014), eggNOG v. 2.1.0 (Huerta-Cepas *et al.* 2016), InterProScan v. 5.47-82.0 (Jones *et al.* 2014), and Swiss-Prot (Boutet *et al.* 2016) by diamond BLASTP v. 2.0.8 (Buchfink *et al.* 2015). Additionally, Phobius v. 1.01 (Käll *et al.* 2004) was used to predict transmembrane proteins and SignalP v. 5.0b (Armenteros *et al.* 2019) was used to predict secreted proteins. Problematic gene models flagged by Funannotate were manually curated as needed. To investigate gene model support, we used STAR v. 2.7.5a to align transcriptome reads to the assembly and then used featureCounts v1.6.2 to generate read counts per gene model (Dobin *et al.* 2013; Liao *et al.* 2014). Read counts per gene model were then summarized in R v. 4.0.3 using the tidyverse v. 1.3.0 package (Wickham *et al.* 2019; R Core Team 2020).

### Identification of genes of interest for future experimental work

We identified candidate genes that could be used as either (1) phenotypic markers, or (2) whose promoters may prove useful for future manipulative experiments using CRISPR technologies. Protein sequences for genes of interest were identified and downloaded from a variety of sources including (1) FlyBase (https://flybase.org/) to obtain orthologs in *Drosophila melanogaster,* (2) FlyBase to identify the closest Hemiptera annotated orthologs, and (3) the literature (Table S2). Protein sequences were searched against the draft *H. vitripennis* genome using phmmer in HMMER v. 3.3.1 (Eddy 2011). Top hits were aligned using MUSCLE v. 3.8.1551 (Edgar 2004). Maximum likelihood trees based on these alignments were produced using FastTree v. 2.0.0 (Price *et al.* 2010) to confirm putative candidate status.

## Results and Discussion

### H. vitripennis predicted genome characteristics

Genome size estimates from GenomeScope ranged from 1.74 to 1.75 Gb, whereas estimates from findGSE were higher, ranging from 1.89 to 1.96 Gb (Figure 1A, Table 1). Both of these approximations are larger than the size of the i5k project reference assembly (1.44 GB). Despite the diversity represented by the Hemiptera (~82,000 species), relatively few genome sequences for this group are available (Panfilio and Angelini 2018). The predicted genome size of *H. vitripennis* fits within the current reported range of genome size estimates for Hemiptera (from ~327 Mb in aphids (Biello *et al.* 2021) to ~8.9 Gb in spittlebugs (Rodrigues *et al.* 2016)) with bloated genome sizes predicted for many members of the Auchenorrhyncha, particularly members of the Cicadidae (Hanrahan and Johnston 2011; Panfilio and Angelini 2018). The predicted heterozygosity of the assembly here was high with the GenomeScope estimates ranging from 1.56 to 1.68%, while the findGSE estimates were lower, ranging from 1.16 to 1.29%. High heterozygosity is not uncommon in Hemiptera and has been reported in planthoppers (Zhu *et al.* 2017), milkweed bugs (Panfilio *et al.* 2019), and aphids (Mathers *et al.* 2020).

**Figure 1.**
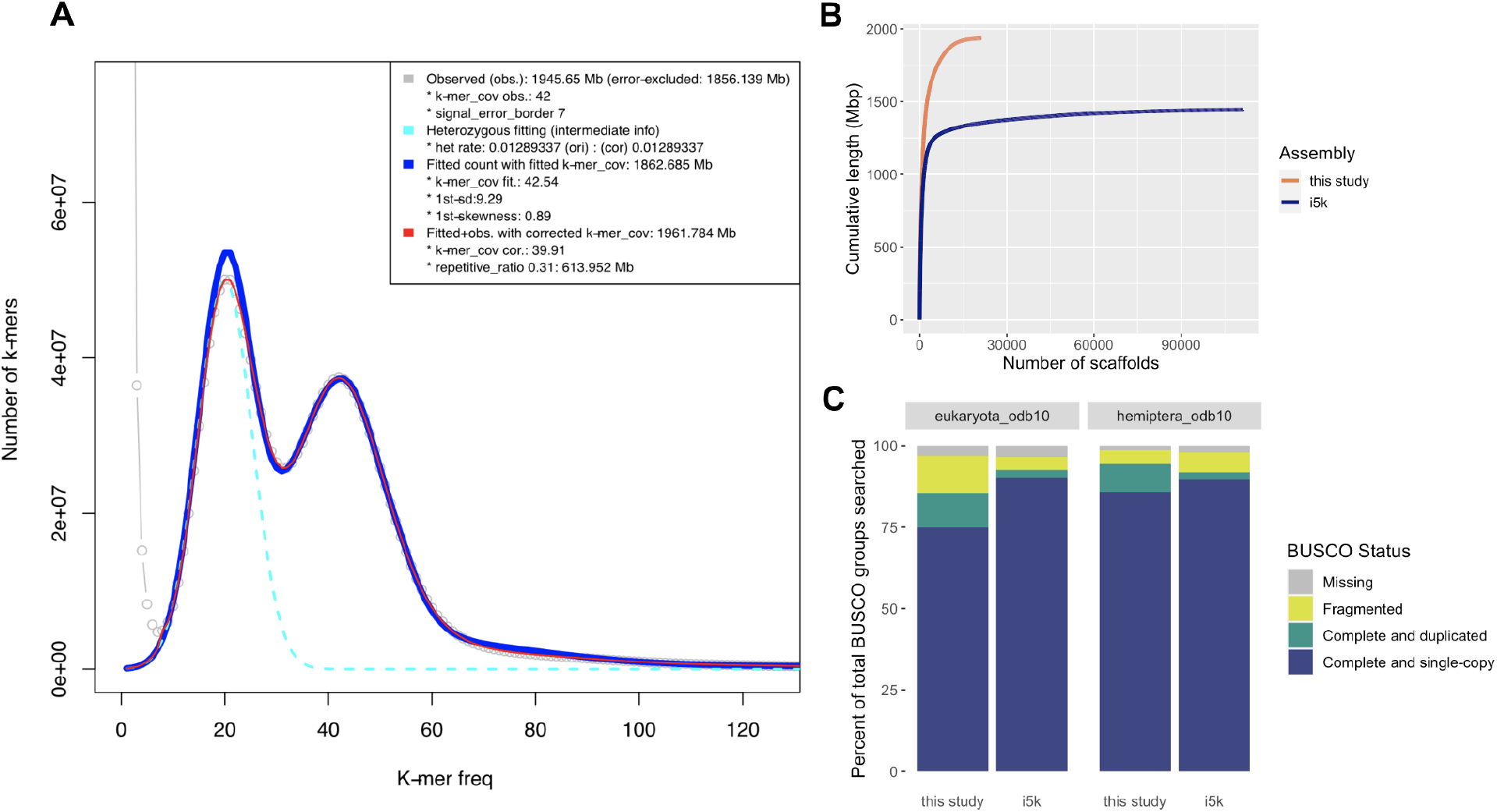
Genome assembly assessment and comparison. (A) *k-*mer frequency histogram output from findGSE using k = 21. The grey line represents the observed *k*-mer frequency, the teal line represents the fit for the heterozygous k-mer peak, the blue line represents the fitted model without k-mer correction, and the red line represents the fitted model with k-mer correction, which is used to estimate the genome size. (B) Plot depicting cumulative sequence length (Y axis) as the number of scaffolds increases (X axis) comparing the *H. vitripennis* draft genome in this study to the reference genome from the i5k project. (C) Stacked barcharts depicting BUSCO analyses for the eukarytota_odb10 and hemiptera_odb10 gene sets for both the *H. vitripennis* genome reported here and the i5k reference genome. Bars show the percent of genes found in each assembly as a percentage of the total gene set and are colored by BUSCO status (missing = grey, fragmented = yellow, complete and duplicated = green, complete and single-copy = blue).

**Table 1.** Estimates of genome heterozygosity, length, and repeat content. These estimates include the percentage heterozygosity, haploid genome size and percentage of repeat content based on *k*-mer analysis using GenomeScope and findGSE for a range of *k*-mers (k = 19, 21, 23, 25, 27).

|  |  | Genomescope | findGSE |
| --- | --- | --- | --- |
| $k = 19$ | Heterozygosity (%) | 1.65 | 1.26 |
|  | Genome haploid size (Gb) | 1.74 | 1.9 |
|  | Repeat (%) | 45.86 | 37.79 |
| $k = 21$ | Heterozygosity (%) | 1.68 | 1.29 |
|  | Genome haploid size (Gb) | 1.74 | 1.96 |
|  | Repeat (%) | 36.91 | 31.3 |
| $k = 23$ | Heterozygosity (%) | 1.65 | 1.29 |
|  | Genome haploid size (Gb) | 1.75 | 1.93 |
|  | Repeat (%) | 34.28 | 28.93 |
| $k = 25$ | Heterozygosity (%) | 1.6 | 1.27 |
|  | Genome haploid size (Gb) | 1.75 | 1.89 |
|  | Repeat (%) | 33.14 | 27.55 |
| $k = 27$ | Heterozygosity (%) | 1.56 | 1.16 |
|  | Genome haploid size (Gb) | 1.75 | 1.96 |
|  | Repeat (%) | 32.37 | 27.03 |

### H. vitripennis genome assembly

The resulting *H. vitripennis* draft genome was assembled into 21,254 scaffolds totalling 1.93 Gb of sequence at 71x coverage with an N50 of 650 Mb (Figure 1B, Table 2). This is an improvement over the current i5k project reference genome, which has 111,110 scaffolds and has a similar N50 of 656 Mb. Additionally, the genome length of the assembly here is in-line with the estimated size range from findGSE (1.89-1.96 Gb).

**Table 2.** Assembly statistics and assessment. Various statistics calculated by QUAST for the assembly in this study and the i5k reference assembly are provided here including the number of contigs in the assembly, the number of scaffolds of various lengths, the total assembly length, percent GC, the N50 and the L50. All statistics from QUAST are based on contigs of size >= 3000 bp, unless specifically noted (e.g., “# contigs (>= 0 bp)” and “Total length (>= 0 bp)” include all contigs in each assembly). We also report here the results of the BUSCO assessment of both assemblies using the hemiptera_odb10 and eukaryota_odb10 gene sets.

|  | Assembly | This study | i5k |
| --- | --- | --- | --- |
| **QUAST** | **#contigs** | 34952 | 149799 |
|  | **# scaffolds (≥ 0 bp)** | 21254 | 111110 |
|  | **# scaffolds (≥ 1000 bp)** | 19715 | 59570 |
|  | **# scaffolds (≥ 5000 bp)** | 14959 | 13241 |
|  | **# scaffolds (≥ 10000 bp)** | 12524 | 7359 |
|  | **# scaffolds (≥ 25000 bp)** | 8796 | 4438 |
|  | **# scaffolds (≥ 50000 bp)** | 5168 | 3132 |
|  | **Total length (≥ 0 bp)** | 1930946379 | 1445215006 |
|  | **Total length (≥ 1000 bp)** | 1929918132 | 1418424409 |
|  | **Total length (≥ 5000 bp)** | 1916091697 | 1325420810 |
|  | **Total length (≥ 10000 bp)** | 1898148486 | 1285066097 |
|  | **Total length (≥ 25000 bp)** | 1833358540 | 1240043308 |
|  | **Total length (≥ 50000 bp)** | 1703319989 | 1194181890 |
|  | **Largest contig** | 7378560 | 7131305 |
|  | **GC (%)** | 32.87 | 32.65 |
|  | **N50** | 650435 | 656130 |
|  | **N75** | 171660 | 211051 |
|  | **L50** | 750 | 542 |
|  | **L75** | 2178 | 1423 |
|  | **# N's per 100 kbp** | 71.13 | 3005.46 |
| **hemiptera_odb10** | **Complete BUSCOs (C)** | 2367 (94.3%) | 2306 (91.9%) |
|  | **Complete and single-copy BUSCOs (S)** | 2152 (85.7%) | 2247 (89.5%) |
|  | **Complete and duplicated BUSCOs (D)** | 215 (8.6%) | 59 (2.4%) |
|  | **Fragmented BUSCOs (F)** | 108 (4.3%) | 150 (6.0%) |
|  | **Missing BUSCOs (M)** | 35 (1.4%) | 54 (2.1%) |
|  | **Total BUSCO groups searched** | 2510 | 2510 |
| **eukaryota_odb10** | **Complete BUSCOs (C)** | 218 (85.5%) | 236 (92.6%) |
|  | **Complete and single-copy BUSCOs (S)** | 191 (74.9%) | 230 (90.2%) |
|  | **Complete and duplicated BUSCOs (D)** | 27 (10.6%) | 6 (2.4%) |
|  | **Fragmented BUSCOs (F)** | 29 (11.4%) | 10 (3.9%) |
|  | **Missing BUSCOs (M)** | 8 (3.1%) | 9 (3.5%) |
|  | **Total BUSCO groups searched** | 255 | 255 |

Assessment with the BUSCO Hemiptera set showed minor improvement in genome completion (94.3%) over the i5k reference genome (91.9%), but more duplications (8.6% vs. 2.4%) (Figure 1C, Table 2). Using the BUSCO eukaryota_odb10 set, the draft genome here was actually less complete (85.5%) compared to the i5k reference genome (92.6%), although both were similarly complete when taking into account fragmented BUSCOs (96.9% here vs. 96.5% i5k). The increased number of fragmented and duplicated BUSCOs may be in part due to the moderate heterozygosity (e.g. haplotigs - allelic variants assembled as separate scaffolds), but is more likely the result of poor assembly of repetitive regions given the relatively high proportion of genome predicted to be repetitive (described below).

### Endosymbiont identification and assessment includes high-quality draft MAG from Wolbachia sp

Like many sap-feeding insects, *H. vitripennis* relies on obligate symbioses with bacterial species for biosynthesis of essential amino acids, which are limited in its xylem-based diet (Wu *et al.* 2006; McCutcheon and Moran 2007). The first of these obligate endosymbionts is *Candidatus* Sulcia muelleri, which has a reduced genome (~243 kb) (Moran *et al.* 2005; Wu *et al.* 2006; McCutcheon *et al.* 2009). The second obligate endosymbiont is *Ca*. Baumannia cicadellinicola has a relatively larger genome (~686 kb) likely due to its more recent acquisition by *H. vitripennis* as a symbiont (Moran *et al.* 2003; Wu *et al.* 2006; Bennett and Moran 2013; Moran and Bennett 2014). In addition to these two obligate symbionts, *Wolbachia* sp. have been observed as abundant facultative symbionts in this species (Moran *et al.* 2003; Wu *et al.* 2006; Curley *et al.* 2007; Hail *et al.* 2011; Rogers and Backus 2014; Welch *et al.* 2015; Pascar and Chandler 2018).

To identify potential contaminant reads due to obligate or facultative symbionts in the *H. vitripennis* draft genome, we used two complementary methods, BlobTools2 and anvi’o (Figure 2). BlobTools2 flagged 167 scaffolds as possible contaminants (Figure 2A). Of these, 19 were confirmed to also belong to draft MAGs assembled in anvi’o, and all scaffolds mapping to MAGs were subsequently removed from the *H. vitripennis* assembly. In total, we generated four draft MAGs for removal from the *H. vitripennis* draft genome assembly (Table 3). These included one near-complete (> 99%) high-quality *Wolbachia* sp. MAG (Figure 2B), one partial *Ca.* Baumannia cicadellinicola MAG (Figure 2C), and two partial *Ca.* Sulcia muelleri MAGs (Figure 2D). Genomic comparisons between *Wolbachia* sp. (GWSS-01) and other *Wolbachia* sp. may help shed light on the possible function (or lack thereof) of this facultative endosymbiont when associated with *H. vitripennis*. Additionally, we hope that this MAG may serve as a useful resource for potential *Wolbachia*-mediated insect-control for *H. vitripennis* in its invasive range (Zabalou *et al.* 2004; Brelsfoard and Dobson 2009; Bourtzis *et al.* 2014).

**Figure 2.**
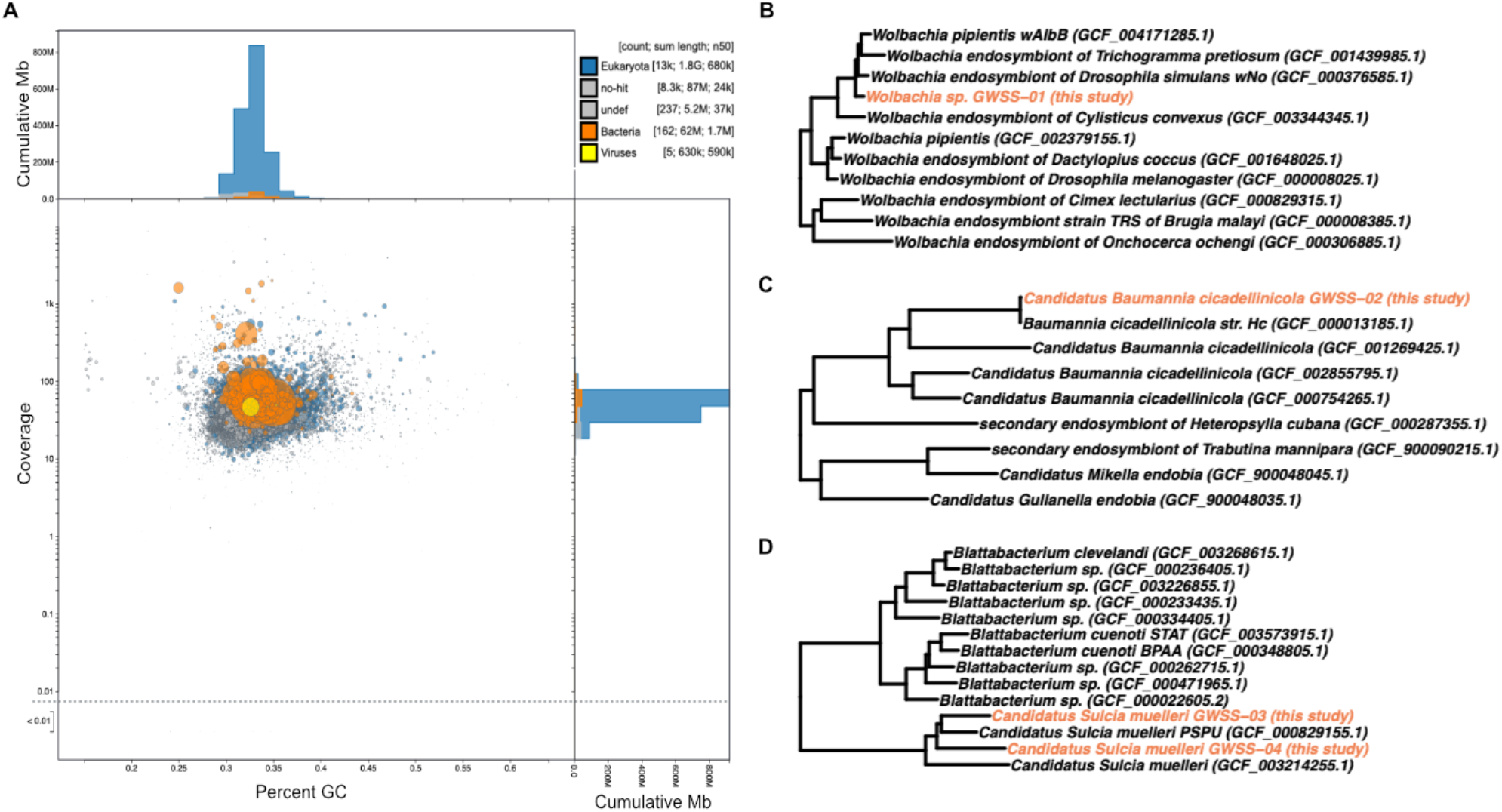
Endosymbiont assessment in genome and identification. (A) BlobTools2 visualization of *H. vitripennis* scaffolds showing taxa-colored GC coverage plot. Each circle represents a scaffold in the assembly, scaled by length, and colored by superkingdom (eukaryota = blue, bacteria = orange, viruses = yellow, unidentified = grey). On the X axis is the average GC content of each scaffold and on the Y axis is the average coverage of each scaffold to the draft assembly. The marginal histograms show cumulative genome length (Mb) for coverage (Y axis) and GC content bins (X axis). (B) Placement of *Wolbachia* sp. GWSS-01 (colored in orange) in the GTDB phylogenetic tree. (C) Placement of *Ca.* Baumannia cicadellinicola GWSS-02 (colored in orange) in the GTDB phylogenetic tree. (D) Placement of *Ca.* Sulcia muelleri GWSS-03 and GWSS-04 (colored in orange) in the GTDB phylogenetic tree.

**Table 3.** Genome feature summary for endosymbiont MAGs. Genomic characteristics are summarized for each metagenome-assembled genome (MAG), including putative taxonomic identity, length (bp), number of scaffolds, N50, percent GC content, number of genes, presence of 16S ribosomal RNA gene, and completion and contamination estimates as generated by CheckM. MAGs are sorted by percent completion.

| MAG ID | Taxonomy | Total length (bp) | Number of scaffolds | N50 | GC (%) | Number of genes | 16S rRNA copy present | Completion (%) | Redundancy (%) |
| --- | --- | --- | --- | --- | --- | --- | --- | --- | --- |
| GWSS-01 | *Wolbachia sp.* | 1712771 | 1 | 1712771 | 33.66 | 1,691 | yes | 99.36 | 1.71 |
| GWSS-02 | *Ca. Baumannia cicadellinicola* | 610888 | 12 | 78712 | 32.65 | 531 | yes | 66.46 | 1.25 |
| GWSS-03 | *Ca. Sulcia muelleri* | 209259 | 1 | 209259 | 24.95 | 199 | yes | 25.86 | 0 |
| GWSS-04 | *Ca. Sulcia muelleri* | 179112 | 6 | 41952 | 26.84 | 148 | no | 17.76 | 1.34 |

### H. vitripennis genome annotation

In total, 98,296 protein-coding genes (91.5% of which are complete with both a stop and start codon) and 10,466 tRNA genes were predicted in the *H. vitripennis* using the funannotate pipeline (Table 4). This is almost twice the number reported by the previous transcriptome effort (47,265 protein-coding genes) (Nandety *et al.* 2013), but is consistent with the number of transcripts (106,998) reported for the transcriptome of *H. liturata* (Tassone *et al.* 2017). Of the 98,296 protein-coding genes reported here, approximately 38.3% (37,652) had at least one database match. In comparison, 45% (23,547) of predicted proteins in the transcriptome reported by Nandety et al (2013) had database matches. The mean annotation edit distances (AED) reported for the predicted coding sequences (CDS) was 0.002 and for the mRNA was 0.024.

**Table 4.**
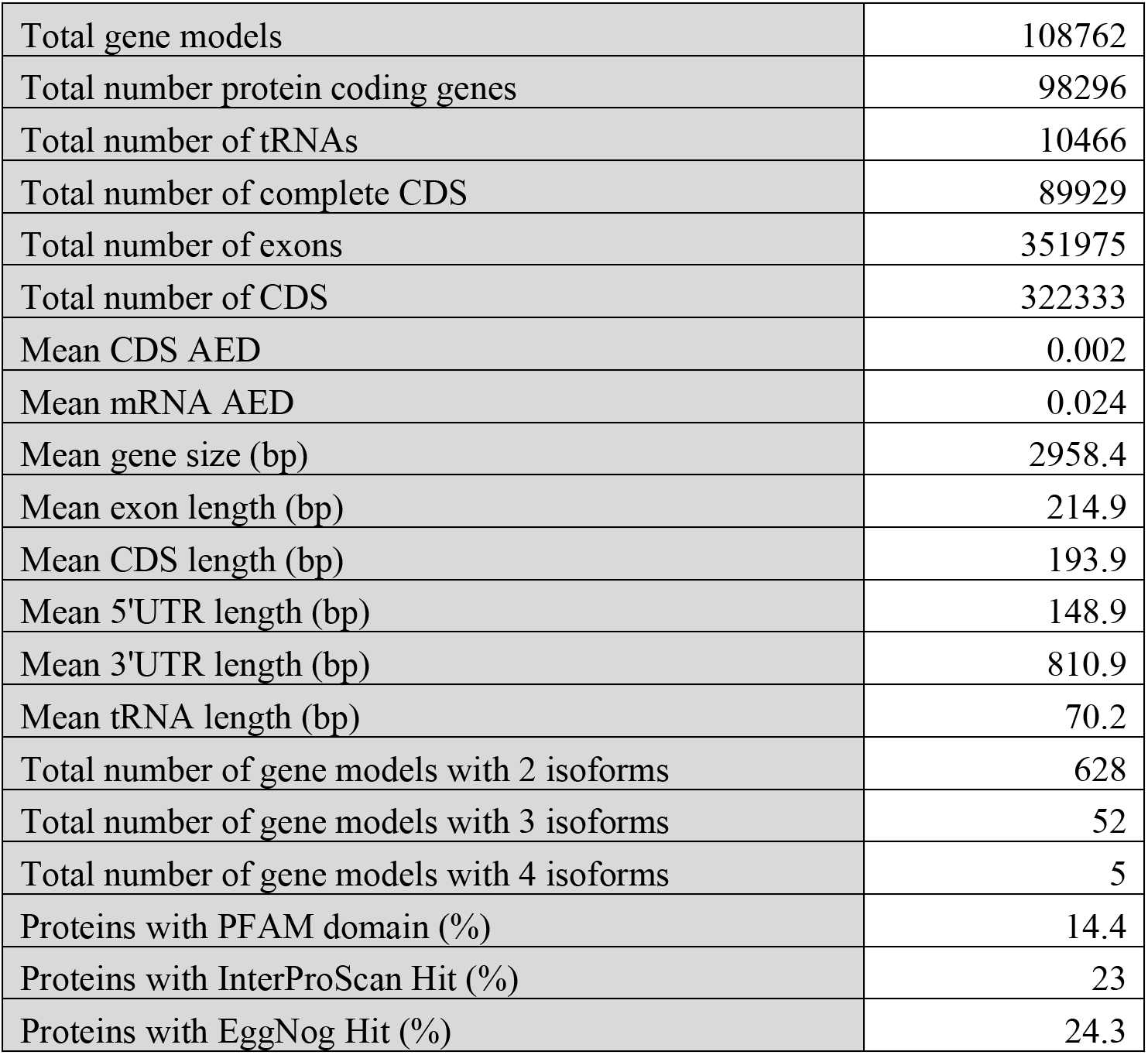
Genome annotation statistics. A summary of genome annotation results is reported here including the total number of gene models, protein-coding genes, tRNAs, complete (e.g. having both a start and stop codon) coding sequences (CDS), exons, and CDS regions, the mean CDS and mRNA annotation edit distances (AED), the mean gene size (bp), exon length (bp), CDS length (bp), 5’-UTR length (bp), 3’-UTR length (bp), and tRNA length (bp), the total number of gene models with 2, 3 or 4 isoforms, and the percentage of proteins with a PFAM domain, InterProScan or EggNog match.

AEDs are a measure of concordance between the gene models and input evidence (such as the transcriptome evidence provided to PASA) with low values like those obtained in this study indicating support for gene models. Further, 58.6% (63,358) of gene models had at least one transcriptome read that aligned in our post-annotation assessment, indicating strong support for at least half of the predicted models. Only adult prothoracic leg tissue transcriptomes were sequenced here, so this value is likely an underestimate. Additional transcriptome data across a range of body parts and developmental stages would be necessary to further confirm the remaining predictions.

The number of protein-coding genes predicted here, although similar to the number reported from the transcriptome of *H. liturata,* is substantially higher than the number of curated predictions from genomes of other Hemiptera species (ranging from 15,456 in *Rhodnius prolixus* (Mesquita et al. 2015) to 36,985 in *Cimex lectularius* (Rosenfeld et al. 2016)). However, as of 2019, only 16 curated genome annotations were available in NCBI belonging to members of Hemiptera (Li *et al.* 2019). With the lack of reference genomes and annotations for Hemiptera, additional sequencing and annotation of close relatives of *H. vitripennis* may reveal similarly increased numbers of gene models. Given the high heterozygosity of the genome, however, overestimation or fragmentation of gene models during the predictions cannot be completely ruled out. Leveraging long-read sequencing technologies should help to further overcome any remaining gene-model fragmentation and future work should seek to validate and refine these predicted gene models.

### Repeat landscape indicates two possible expansion events

The estimated percentage of the genome that was repetitive was relatively high (GenomeScope = 32.37-45.86%; findGSE = 27.03-37.79%), with ultimately 33.06% of the genome being identified and masked as repetitive by RepeatMasker (see Table S1 for detailed breakdown). This value is similar to other Hemiptera genomes, e.g. 23.0% in *Laodelphax striatellus* (Zhu *et al.* 2017), 38.9% in *Nilaparvata lugens* (Xue *et al.* 2014), 39.7% in *Sogatella furcifera* (Zhu *et al.* 2017), 45% in *Bemisia tabaci* (Chen et al. 2016), 56.6% in *Trialeurodes vaporariorum* (Xie et al. 2020), and 60% in *Locusta migratoria* (Wang *et al.* 2014), as well as other insect genomes, e.g. 33% in *Tribolium castaneum* (Tribolium Genome Sequencing Consortium *et al.* 2008), 40% in *Bombyx mori* (Cai *et al.* 2012), and 47% in *Aedes aegypti* (Nene *et al.* 2007). However, in contrast, Nandety et al. (2013) reported that only ~1% of the *H. vitripennis* transcriptome represented repetitive elements. One possible explanation for this is that the majority of repeat content in *H. vitripennis* is not in coding regions and was not captured by previous transcriptome efforts.

The most abundant repeat elements (~18% of genome) in *H. vitripennis* were unclassified, followed by LINES (~6.7% of genome) and DNA elements (5.8% of genome) (Figure 3A, Table S1). Generally, this repeat element diversity was consistent with other Hemiptera (e.g. (Petersen *et al.* 2019)). Additionally, the repeat landscape indicates that elements have accumulated gradually through time in this species and also exposes two possible expansions of repeat content, one ancient (corresponding to ~21% divergence) and one more recent (corresponding to 2-4% divergence) (Figure 3B).

**Figure 3.**
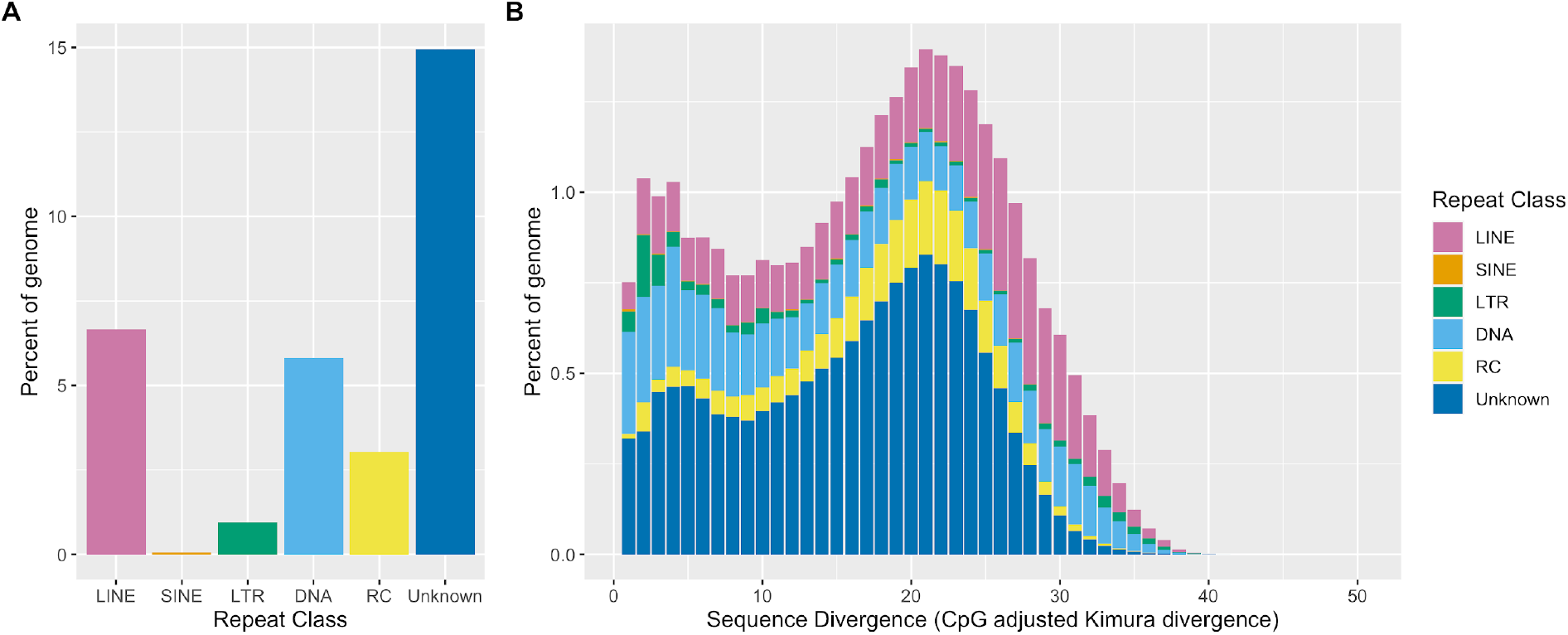
Repetitive element diversity and divergence landscape. (A) A barplot representing the percent of the genome composed of elements from each repeat class. (B) A stacked barplot representing the percent of the genome made of repeat elements from each repeat class binned by 1% sequence divergence (CpG adjusted Kimura divergence). Bars are colored repeat class (LINE = pink, SINE = orange, LTR = green, DNA = light blue, RC = yellow, Unknown = dark blue). Abbreviations: long-interspersed nuclear element (LINE), small-interspersed nuclear element (SINE), long-terminal repeat retrotransposon (LTR), DNA transposons (DNA), rolling-circle transposons (RC).

### Identification of 27 candidate genes as tools for use in genetic analyses

We identified 14 candidate genes that can be used as phenotypic markers and 13 candidate genes whose promoters may prove useful for future manipulative experiments (e.g using CRISPR technologies) (Table 5). Of the 14 candidate morphological markers identified, nine are involved in eye color, one in body color, three in wing morphology, and one in eye morphology. These phenotypes are predicted based on known phenotypes in *Drosophila melanogaster* where these genes have been useful resources for genetic analysis for years (Chyb and Gompel 2013). For the 13 candidate genes with promoters of interest, we searched for and identified four *actin* genes, two *polyubiquitin* genes, one *exuperantia* (*exu*) gene, one *vasa* gene and five *beta-tubulin* genes. In order to genetically manipulate *H. vitripennis*, we first need to identify genes with promoters that are constitutively expressed, or expressed in tissues and developmental stages of interest. We believe that the reported collection of phenotypic marker genes and genes with promoters of interest, will be a useful resource for the community of researchers using genetic tools in this species.

**Table 5.**
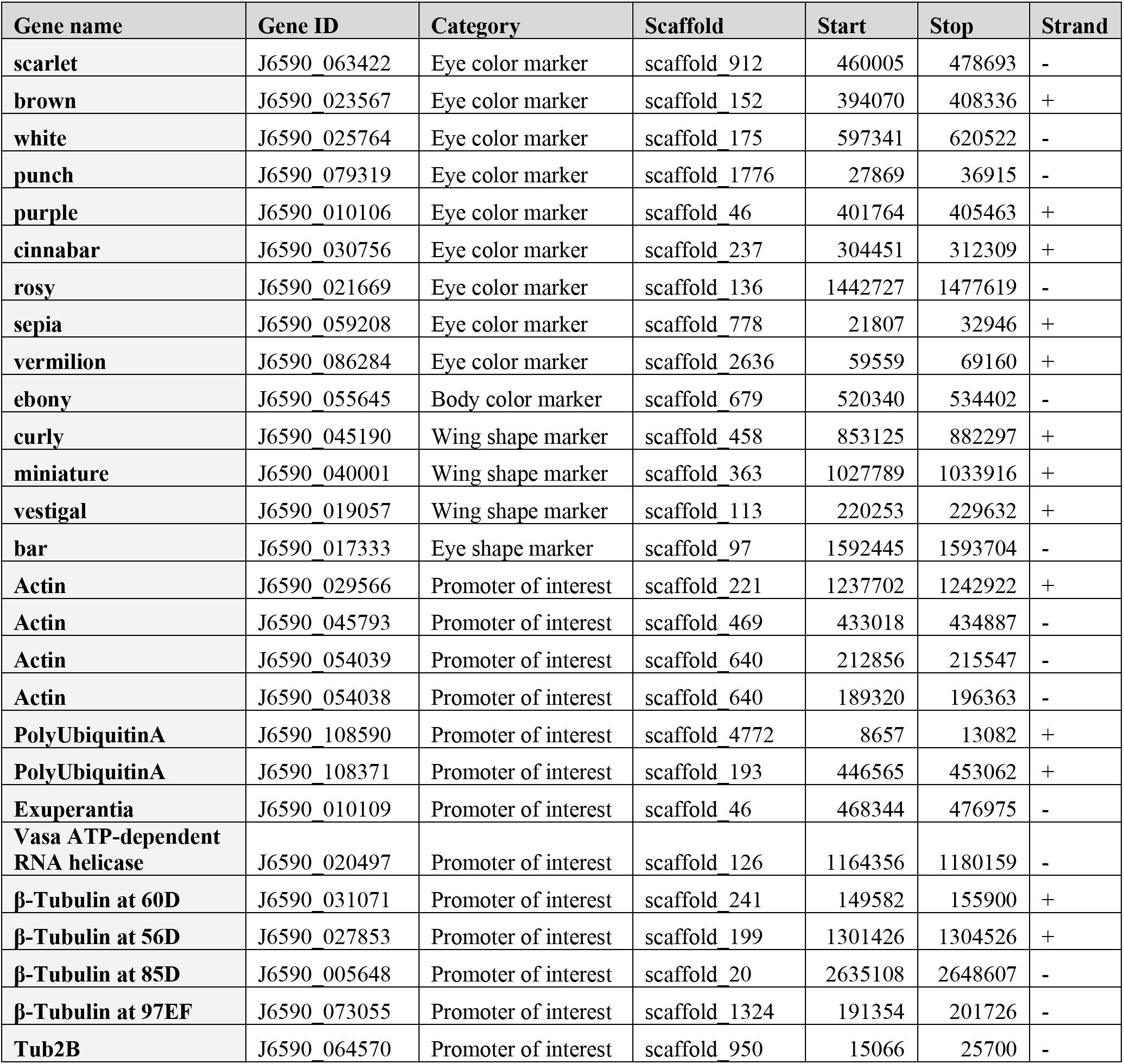
Orthologous candidate genes identified for use in genetic analyses. Here for each identified gene, we provide the gene name, gene ID (e.g. the loci name provided to NCBI), scaffold number, strand direction, and start and stop locations. We also report the category of interest for each gene. Broadly, these fall into two larger groupings: (1) promoter of interest or (2) a morphological marker category based on phenotype from the literature (e.g. eye color, body color, wing shape, eye shape).

## Conclusions

Using a combination of Oxford Nanopore long-read and Illumina short-read technologies, we generated an improved reference genome for *H. vitripennis* of 21,254 scaffolds and a total genome size of 1.93 Gb. As part of this process, we also assembled four endosymbiont genomes, including a high-quality near complete *Wolbachia* sp. We further provide a first pass at genome annotation for *H. vitripennis*, predicting 98,296 protein-coding genes and 10,466 tRNA genes, of which 38.3% had homology matches to current databases. As an additional community resource, we identified 27 orthologous candidate genes of interest to be leveraged in future studies that seek to genetically manipulate *H. vitripennis*. Given the increasing role of *H. vitripennis* as an invasive agricultural pest, we hope that the generated genome assembly, endosymbiont MAGs, annotation and curated set of candidate genes will serve as important resources for future genomics, genetics, biocontrol and insect biology research of *H. vitripennis,* other sharpshooters, and leafhoppers.

## Data Availability

The draft *H. vitripennis* Tulare genome assembly, annotation, and mitochondrial genome are deposited at DDBJ/ENA/GenBank under the accession JAGXCG000000000. The version described in this paper is version JAGXCG010000000. The raw sequence reads for the genome and RNA-Seq are available through BioProjects PRJNA717305 and PRJNA717315, respectively. The four MAG assemblies are available from BioProject PRJNA723626 and are deposited at DDBJ/ENA/GenBank under accession numbers JAGTUP000000000, JAGTUQ000000000, JAGTUR000000000, and JAGTUS000000000. Data analysis, assembly and annotation related scripts for this work are available on GitHub and archived in Zenodo (Ettinger and Stajich 2021).

## Acknowledgments

JES is a CIFAR Fellow in the program Fungal Kingdom: Threats and Opportunities and partially supported by USDA Agriculture Experimental Station at the University of California, Riverside and NIFA Hatch projects CA-R-PPA-5062-H. This work was supported by the Pierce’s Disease Control program (sponsor award #: 14-0379-000-SA-2) to FJB and RR and California Department of Food and Agriculture agreement 20-0267 to RR, LLW, PWA and JES.

## Notes

### Competing Interest Statement

The authors have declared no competing interest.

