## Supplemental Table Legends for "Improved draft reference genome for the Glassy-winged Sharpshooter (*Homalodisca vitripennis*), a vector for Pierce’s disease"

**Table S1. Repeat content summary.** Repeat content results are reported here including the percentage of the genome identified and masked per type repeat element by RepeatMasker. Abbreviations of repeat element categories: long-interspersed nuclear element (LINE), small-interspersed nuclear element (SINE), long-terminal repeat retrotransposon (LTR), DNA transposons (DNA), and rolling-circle transposons (RC).

**Table S2. Reference alleles used to identify candidate genes.** For each candidate gene, we provide the gene name, the FlyBase ID (*Drosophila melanogaster,* <https://flybase.org/>) for the gene used, an alternative insect species ortholog ID used and the species the alternate ID represents. We also provide the category of interest for each gene. Broadly, these fall into two larger groupings: (1) promoter of interest or (2) a morphological marker category based on phenotype from the literature (e.g. eye color, body color, wing shape, eye shape).
